# Genetically encoded tool for manipulation of ΔΨm identifies the latter as the driver of integrative stress response induced by ATP Synthase dysfunction

**DOI:** 10.1101/2023.12.27.573435

**Authors:** Mangyu Choe, Denis V. Titov

## Abstract

Mitochondrial membrane potential (ΔΨm) is one of the key parameters controlling cellular bioenergetics. Investigation of the role of ΔΨm in live cells is complicated by a lack of tools for its direct manipulation without off-target effects. Here, we adopted the uncoupling protein UCP1 from brown adipocytes as a genetically encoded tool for direct manipulation of ΔΨm. We validated the ability of exogenously expressed UCP1 to induce uncoupled respiration and lower ΔΨm in mammalian cells. UCP1 expression lowered ΔΨm to the same extent as chemical uncouplers but did not inhibit cell proliferation, suggesting that it manipulates ΔΨm without the off-target effects of chemical uncouplers. Using UCP1, we revealed that elevated ΔΨm is the driver of the Integrated Stress Response induced by ATP synthase inhibition in mammalian cells.

## Introduction

Mitochondrial membrane potential (ΔΨm) is an important regulator of mitochondrial function. ΔΨm of 100 to 150 mV is generated by proton pumps of the mitochondrial electron transport chain (ETC)–Complexes I, III and IV. The energy conserved in ΔΨm is, in turn, used for ATP production by mitochondrial ATP synthase or Complex V. In addition to the canonical role in ATP production, ΔΨm also regulates metabolite transport, protein import, ROS production^1–4^. Furthermore, ΔΨm plays a role in the quality control machinery of mitochondria, enabling the disposal of dysfunctional mitochondria, and has been implicated as a modulator of aging^5–8^.

A key barrier to a better understanding of the function of ΔΨm is the lack of tools for direct and specific manipulation of ΔΨm. Several reagents have been developed to target ΔΨm, including chemical uncouplers (i.e., Carbonyl cyanide-p-trifuloremethoxyphenylhydrazone (FCCP), Bam15, and 2,4-dinitrophenol (DNP)) and optogenetic tools (mt^SWITCH^)^9–12^. Current approaches to manipulate ΔΨm have several limitations, such as the off-target effects of chemical uncouplers, which uncouple all membrane potentials and not just mitochondrial^13^, and the need for specialized equipment that limits the application of optogenetic tools. Thus, novel tools for manipulating ΔΨm are needed.

In this study, we used UCP1 as a genetically encoded tool to directly manipulate ΔΨm. UCP1 is predominantly expressed in brown adipocytes, where it dissipates ΔΨm to generate heat instead of ATP and is believed to be one of the key mediators of non-shivering thermogenesis^14,15^ (Fig. **1**a). UCP1 dissipates ΔΨm by an unusual mechanism requiring long-chain fatty acids as a catalyst. Long-chain fatty acid binds to UCP1, and the carboxylic group of the fatty acid swings back and forth across the mitochondrial inner membrane to transport protons down the electrochemical gradient and into the mitochondrial matrix^16,17^. UCP1 has not been employed as a tool to manipulate ΔΨm, but several studies^18,19^ have reported the exogenous expression of active UCP1 in yeast or mammalian cells as part of the effort to better understand if UCP1 alone is sufficient for uncoupled respiration or requires additional partners.

**Fig. 1.**
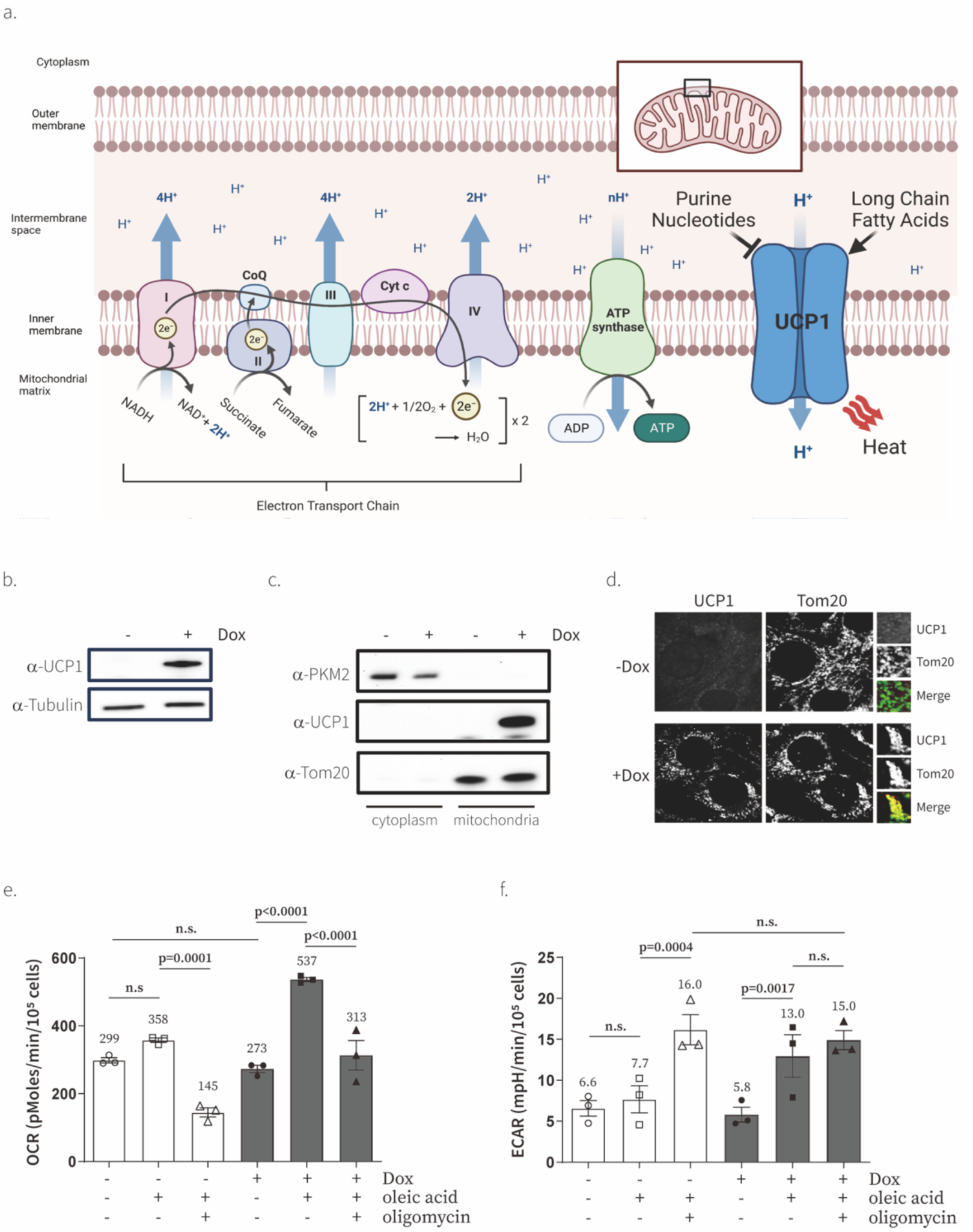
Exogenous expression of UCP1 causes uncoupled respiration in the presence of fatty acids. **a,** A schematic view of the molecular mechanism for the electron transport chain (ETC) and Uncoupling Protein 1 (UCP1). **b**, Western blot of UCP1 in C2C12 cells after 24-hr induction with water or doxycycline (Dox) (300 ng/ml). Representative gel from one of three independent experiments **c**, Subcellular localization of UCP1 in C2C12 cells determined by cell fractionation. PKM2 is a cytosolic marker and Tom20 is a mitochondrial marker. Representative gel from one of two independent experiments. **d**, Subcellular localization of UCP1 in C2C12 cells determined by immunofluorescence. Tom20 is a marker of mitochondria. **e,** Effect of UCP1 expression in C2C12 cells on basal, 300 µM oleic acid-treated, and 300 µM oleic acid & 1 µM oligomycin-treated oxygen consumption rate (OCR) and extracellular acidification rate (ECAR) measured with an XFe24 analyzer. Means ± SEM, n=3 independent experiments. P values were determined with two-way ANOVA followed by Tukey’s multiple comparisons test in Graphpad (n.s., not significant)

Here, we validate UCP1 as a genetically encoded tool for the manipulation of ΔΨm in mammalian cells. We showed that UCP1 specifically dissipates mitochondrial membrane potential without the off-target effects of chemical uncouplers, such as inhibition of cell proliferation. Direct manipulation of ΔΨm rescued the inhibitory effect of oligomycin on cell proliferation. We used UCP1 to demonstrate that elevated ΔΨm drives the Integrated Stress Response and transcriptional response to ATP synthase dysfunction.

## RESULTS

### Exogenous expression and localization of UCP1 in mammalian cells

We used consecutive lentiviral infections to generate C2C12 cells that express mouse UCP1 under the control of doxycycline-inducible promoter TRE3G and doxycycline-sensitive transcription factor Tet3G under the control of EF1alpha promoter. Tet3G induces transcription from the TRE3G promoter in the presence of doxycycline. To increase the expression level of UCP1, we used a strong Kozak sequence (GCCACCATGGGG). C2C12 TRE3G UCP1 cells exhibited doxycycline-inducible UCP1 expression with no detectable leaky expression in the absence of doxycycline (Fig. **1**b). We confirmed the mitochondrial localization of UCP1 using cell fractionation and fluorescence microscopy (Fig. **1**c,d).

### UCP1 causes uncoupled respiration in mammalian cells

To determine whether exogenously expressed UCP1 was active in live cells, we measured its ability to uncouple oxygen consumption from respiratory ATP production. UCP1 uncouples respiration by pumping protons from mitochondrial intermembrane space to the matrix, which leads to the dissipation of ΔΨm^20,21^. The expected result of UCP1 activity would be an increase in oxygen consumption as ETC needs to work faster to pump protons from matrix to intermembrane space to counteract the opposing activity of UCP1, and an increase in lactate secretion as ATP synthase cannot produce ATP as efficiently due to lower ΔΨm. Since catalytic amounts of long-chain fatty acid are required for UCP1 activity^16,17^, we performed our measurements in the presence and absence of physiologically relevant concentrations of oleic acid (300 μM)^22,23^ conjugated to bovine serum albumin (BSA) at 6:1 molar ratio of oleate:BSA^24,25^. We found that UCP1 expression increased oxygen consumption by ∼2-fold in the presence of oleic acid and the UCP1- induced increase in oxygen consumption was resistant to oligomycin treatment as would be predicted for uncoupled respiration (Fig. **1**e and Extended data Fig. 1a). It is noteworthy that UCP1 was inactive in C2C12 cells in the absence of exogenous oleic acid. We also showed that the extracellular acidification rate (ECAR)–a proxy of the fermentative lactic acid production rate–was increased after UCP1 expression (Fig. **1**f and Extended data Fig. 1b). These observations show that exogenously expressed UCP1 is active in the presence of physiologically relevant concentrations of oleic acid and causes uncoupling of oxygen consumption from respiratory ATP production.

### UCP1 rescues oligomycin-induced cell proliferation inhibition

To further validate that exogenously expressed UCP1 is active, we tested its effect on cell proliferation in the presence of ETC inhibitors. Mitochondrial electron transport chain defects lead to the inhibition of mammalian cell proliferation due to a block in NAD^+^ recycling that, in turn, inhibits the biosynthesis of non-essential amino acids^26–28^. This cell proliferation defect can be rescued by stimulating NAD^+^ recycling using LbNOX or pyruvate and by the addition of non-essential amino acid aspartate^26–28^. UCP1 would be predicted to rescue NAD^+^ recycling of ETC in the presence of ATP synthase inhibitors, as its uncoupling activity would stimulate ETC activity, as we showed in Fig. 1f, and Complex I of ETC catalyzes an NAD+ recycling reaction. On the other hand, UCP1 should not rescue cell proliferation in the presence of Complex I, III, or IV inhibitors, as Complex I will be inactive in all the latter conditions. As predicted, we observed that UCP1 could alleviate the inhibitory effect of oligomycin (ATP synthase inhibitor), but not piericidin (Complex I inhibitor), on cell proliferation (Fig. 2a,b,c). We note that, as in the case of uncoupled respiration (Fig. **1**e,f), UCP1 was only active in the presence of exogenous oleic acid. As a positive control, we also showed that UCP1 has no effect in the presence of pyruvate, which could rescue the effect of oligomycin and piericidin on cell proliferation by itself (Fig. 2a,b,c). The ability of UCP1 to rescue oligomycin-induced cell proliferation inhibition provides additional evidence that exogenously expressed UCP1 is active in cells.

**Fig. 2.**
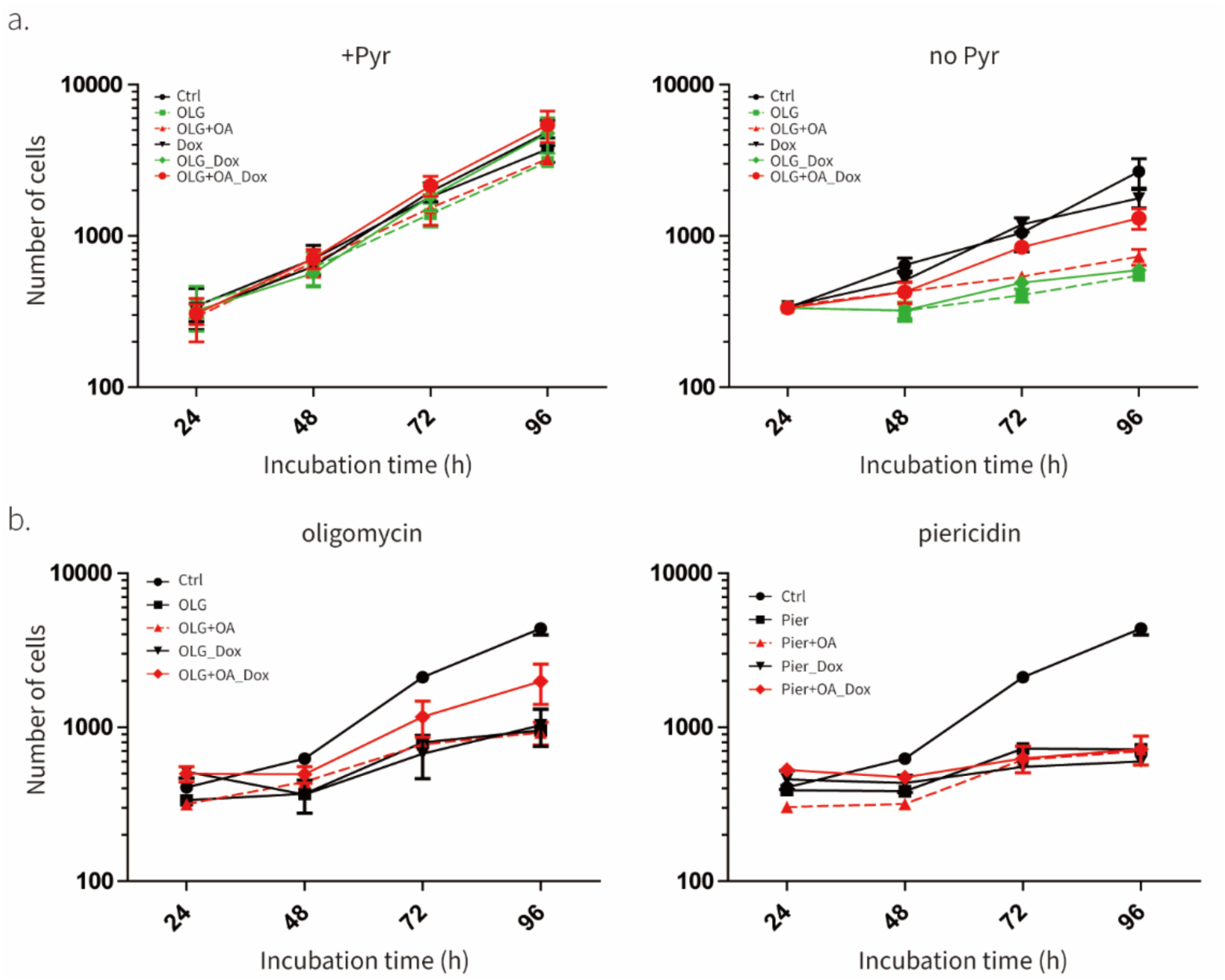
UCP1 rescues the effect of oligomycin on cell proliferation. **a,** Effect of UCP1 expression in C2C12 cells on inhibition of cell proliferation by 1 mM oligomycin (OLG) in the presence (1 mM) or absence of pyruvate. Means ± SEM, n=4 independent experiments. **b,** Effect of UCP1 expression in C2C12 cells on inhibition of cell proliferation by 1 mM oligomycin (OLG) or 1 mM Piericidin (Pier) in the absence of pyruvate. Same data (control) was used in both graphs. Means ± SEM, n=3 independent experiments.

### UCP1 lowers mitochondrial membrane potential

We next tested the ability of UCP1 to lower ΔΨm. To measure ΔΨm, we use TMRM–a cell-permeable, cationic, red-orange fluorescent dye that accumulates inside of the cytoplasm and mitochondria of cells driven by negative plasma and mitochondrial membrane potentials. Thus, TMRM amount inside of cell are high when negative plasma and mitochondrial membrane potentials are high. We validated our TMRM assay by showing that FCCP and Bam15 lower TMRM signal in a dose-dependent manner and oligomycin increases TMRM signal as a result of ΔΨm hyperpolarization in response to ATP synthase inhibition (Fig. 3a). Then we tested the effect of UCP1 on ΔΨm. All experiments were performed in the presence of oleic acid. We observed that UCP1 decreased ΔΨm (Fig. 3b). Treatment with FCCP in the presence of UCP1 could further decrease TMRM signal, likely due to effect of FCCP on plasma membrane^29^. However, treatment with oligomycin could not increase ΔΨm when UCP1 was expressed, suggesting that the effect of UCP1 on ΔΨm is too large to be reversed by ATP synthase inhibition alone. These measurements validate that exogenous UCP1 expression can be used to manipulate ΔΨm in live cells.

**Fig. 3.**
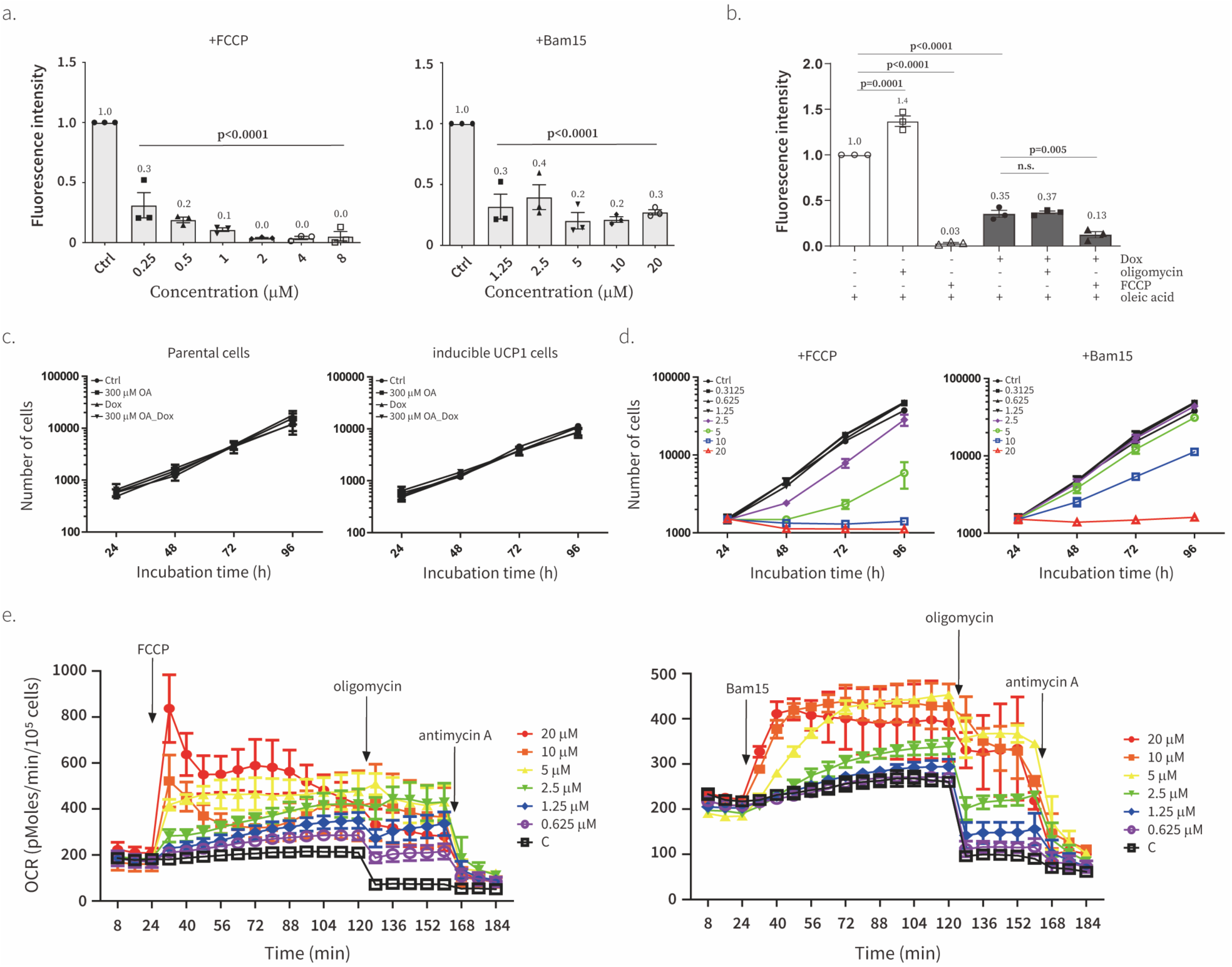
UCP1 lowers ΔΨm without off-target effects of chemical uncouplers. **a**, Effect of various concentrations of FCCP (left) or Bam15 (right) on the ΔΨm using TMRM. The intensity of TMRM was normalized by the intensity of Hoechst. Data is normalized to control condition (Ctrl, lane 1). Means ± SEM, n=3 biological replicates. *P* values were determined with two-way ANOVA followed by Tukey’s multiple comparisons test in Graphpad (*n.s.*, not significant). **b**, Effect of UCP1 on the ΔΨm in the presence of 50 mM oleic acid (Methyl-b-cyclodextrin conjugated) using TMRM. The intensity of TMRM was normalized by the intensity of Hoechst. Data is normalized to oleic acid-treated condition (lane 1). Means ± SEM, n=3 independent experiments. *P* values were determined with two-way ANOVA followed by Tukey’s multiple comparisons test in Graphpad (*n.s.*, not significant). **c**, Effect of oleic acid (OA, 300 mM) or doxycycline (Dox, 300 ng/ml) on proliferation of wild type parental C2C12 (left) or UCP1-expressing (right) C2C12 cells in the presence of 1 mM pyruvate. Means ± SEM, n=3 biological replicates. **d**, Effect of various concentrations of FCCP (left) or Bam15 (right) on the proliferation of C2C12 cells in the presence of 1 mM pyruvate. Means ± SEM, n=4 independent experiments. **e**, Effect of various concentrations of FCCP (left) or Bam15 (right) on basal and oligomycin (1 mM)-treated oxygen consumption. Antimycin A (1 mM) was injected at the last as a negative control. Means ± SEM, n=3 independent experiments.

### UCP1 lowers ΔΨm without the off-target effects of chemical uncouplers

Chemical uncouplers are well known to inhibit cell proliferation^30–33^. Thus, we were surprised when we did not observe the inhibitory effect of UCP1 on cell proliferation (Fig. 2a, Fig 3c), even though the effect of UCP1 on TMRM was similar to that of chemical uncouplers (Fig. 3a,b). Here, we carefully titrated FCCP and Bam15 and compared their effects on uncoupled respiration and cell proliferation to that of UCP1. Optimal concentrations of 5 μM FCCP and 10 μM Bam15 caused a maximal increase in oxygen consumption of about a 2-fold (Fig. 3e), which is similar to the effect of UCP1 (Fig. **1**e). FCCP and Bam15 inhibited cell proliferation at concentrations exceeding 2.5 μM and 5 μM, respectively. Notably, the concentrations of FCCP and Bam15 that start to inhibit cell proliferation are similar or even lower than concentrations causing maximal uncoupled respiration. Comparison of the effect of FCCP, Bam15, and UCP1 on cell proliferation, TMRM signal, and uncoupled respiration suggests that inhibition of cell proliferation constitutes an off-target effect of uncouplers that is unrelated to ΔΨm manipulation, underscoring the importance of exercising caution when attributing the phenotype of an uncoupler treatment to ΔΨm manipulation. Our results suggest that UCP1 is a more specific manipulator of ΔΨm compared to chemical uncouplers and should be preferred to the latter as a reagent to manipulate ΔΨm.

### UCP1 alleviates oligomycin-induced integrative stress response

Having validated UCP1 as tool for targeted manipulation of ΔΨm, we applied it to study the role of ΔΨm in causing the integrative stress response (ISR) in response to mitochondrial dysfunction^34^. ISR is induced by a variety of stressors and the key marker of ISR is the expression of Activating Transcription Factor 4 (ATF4) (**Error! Reference source not found.**Fig. 4a). Most downstream target genes related to ISR are regulated by ATF4. Previously, it was shown that the NADH/NAD^+^ ratio is the signal for ISR after ETC inhibition in proliferating cells, but ATP synthase dysfunction in proliferating and non-proliferating cells induced ISR through a different mechanism^34^. Unlike ETC inhibitors, ATP synthase dysfunction causes elevated ΔΨm in addition to lowered NADH/NAD^+^, implicating ΔΨm as a potential driver of the ISR in response to ATP synthase dysfunction.

**Fig. 4.**
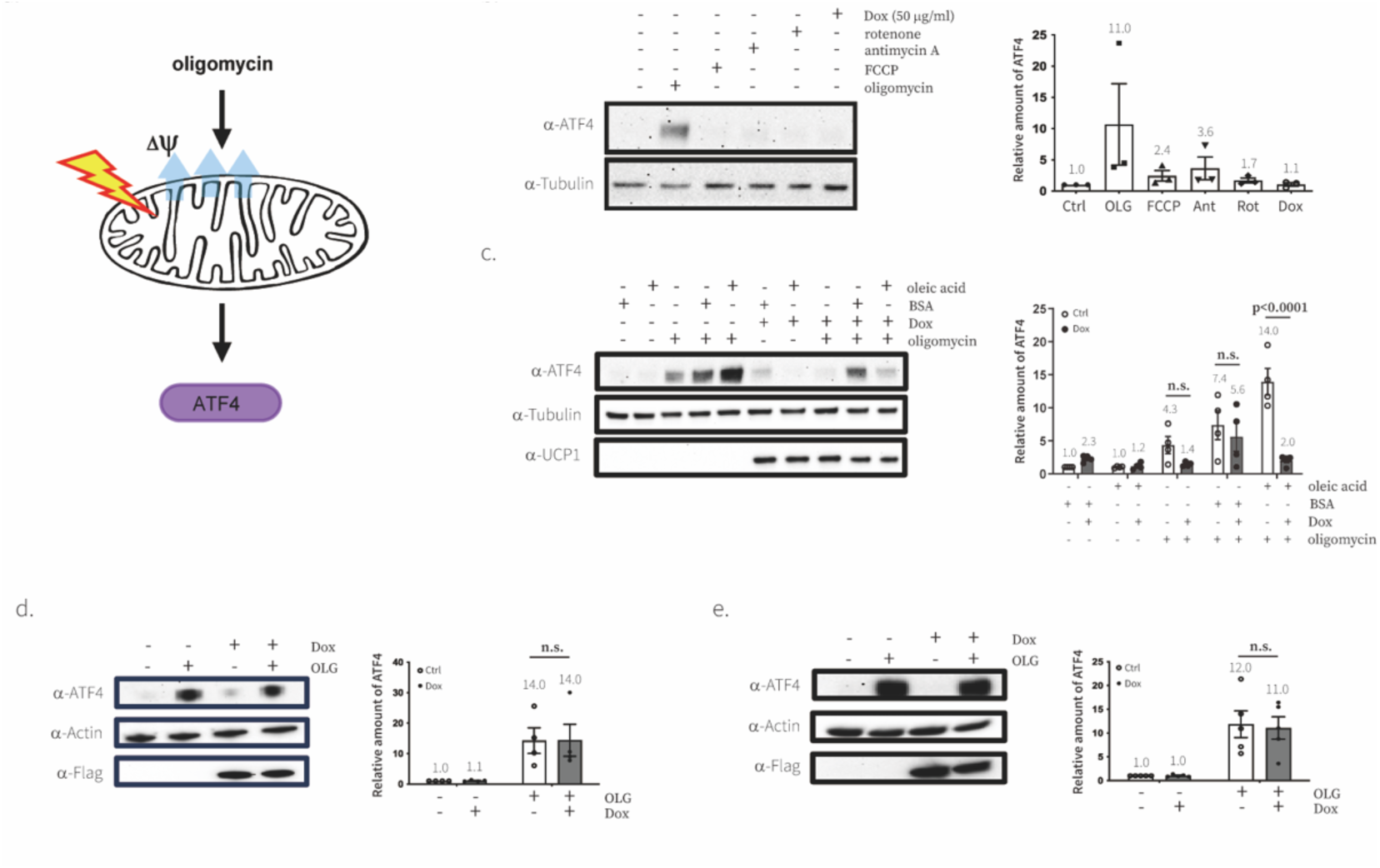
Mitochondrial stress response to ATP synthase inhibition is driven by ΔΨm in C2C12 cells. **a**, A simple schematic diagram showing the molecular mechanism of how oligomycin induces mitochondrial stress signaling. **b**, Effect of various chemical drugs on inducing ATF4 in C2C12 cells. Immunoblotting analyses were performed with the indicated antibodies. Representative gel from one of three experiments. C, control; OLG, 1 mM oligomycin; FCCP, 5 mM FCCP; Ant, 1 mM Antimycin A; Rot, 1 mM Rotenone; Dox, 50 mg/ml doxycycline. Bar graphs show the quantification of the intensity of western blot data, relative amount of ATF4 compared to control condition. Means ± SEM, n=3independent experiments. P values were determined with two-way ANOVA followed by Tukey’s multiple comparisons test in Graphpad (n.s., not significant). **c,d,e**, Effect of UCP1 (c), LbNOX (d), or mitoLbNOX (e) on mitochondrial stress signaling. Representative gel from one of four (UCP1 and LbNOX) or five (mitoLbNOX) experiments Bar graphs show the quantification of the intensity of western blot data, relative amount of ATF4 (left) compared to control condition. Means ± SEM, n=4 or 5 independent experiments. P values were determined with two-way ANOVA followed by Tukey’s multiple comparisons test in Graphpad (n.s., not significant).

Here, we used UCP1 to test if Complex V dysfunction caused by oligomycin treatment induces ISR through elevated ΔΨm. We first confirmed that oligomycin treatment, not other respiration inhibitors, reproducibly and significantly induced the expression of ATF4 even in the presence of pyruvate where the NADH/NAD^+^ ratio is restored as is previously reported (Fig. 4b). We next showed that UCP1 could rescue the effect of oligomycin on ATF4 protein levels (Fig. 4c). We also employed qPCR to demonstrate that ATF4 and GADD34, a representative downstream targets of ATF4, are regulated at the transcription level as well (Extended data Fig. 2). Based on these data, we concluded that elevated mitochondrial membrane potential causes oligomycin-driven mitochondrial stress signaling and direct manipulation of ΔΨm using UCP1 was sufficient to rescue the phenotypes.

### UCP1 rescues oligomycin-driven transcriptional response

To further investigate the effect of direct manipulation of ΔΨm by UCP1, we performed mRNA sequencing in various conditions (Fig. 5a). Since direct manipulation of ΔΨm rescued the effect of oligomycin on ISR, we included oligomycin-treated conditions in our mRNA sequencing experiment. In addition, to verify any effects of the treatment of oleic acid, we also include BSA-treated conditions as negative controls. We performed mRNA sequencing measurement in the presence of pyruvate, which rescues NAD/NAD^+^ imbalance caused by oligomycin, and verified that all treatments showed similar cell growth rates so we could exclude the effect of cell proliferation rate and NADH/NAD^+^ ratio changes (Fig. 5b). The direct manipulation of ΔΨm caused a significant change in the expression of only a few genes indicating that ΔΨm changes do not induce a major transcriptional program under the unperturbed conditions that we tested (Fig. 5c and Extended data Fig. 3a). However, oligomycin treatment and activation of UCP1 under oligomycin treatment showed dramatic changes in both transcriptional profiles and principal component analysis (PCA) (Fig. 5c and Extended data Fig. 3b). We conducted gene ontology enrichment analysis using the Reactome library. The top 10 Reactome groups that exhibited the most significant changes under each condition are illustrated in Extended data Figure 3c. Most of the Reactome groups up-regulated or down-regulated due to oligomycin treatment were down-regulated or up-regulated in response to UCP1 activation under oligomycin-treated conditions. We observed that all genes in the HRI/GCN2-ATF4 group were up-regulated in response to oligomycin treatment but down-regulated when UCP1 was activated under the same oligomycin-treated condition (Extended data Fig. 3c,d).

**Fig. 5.**
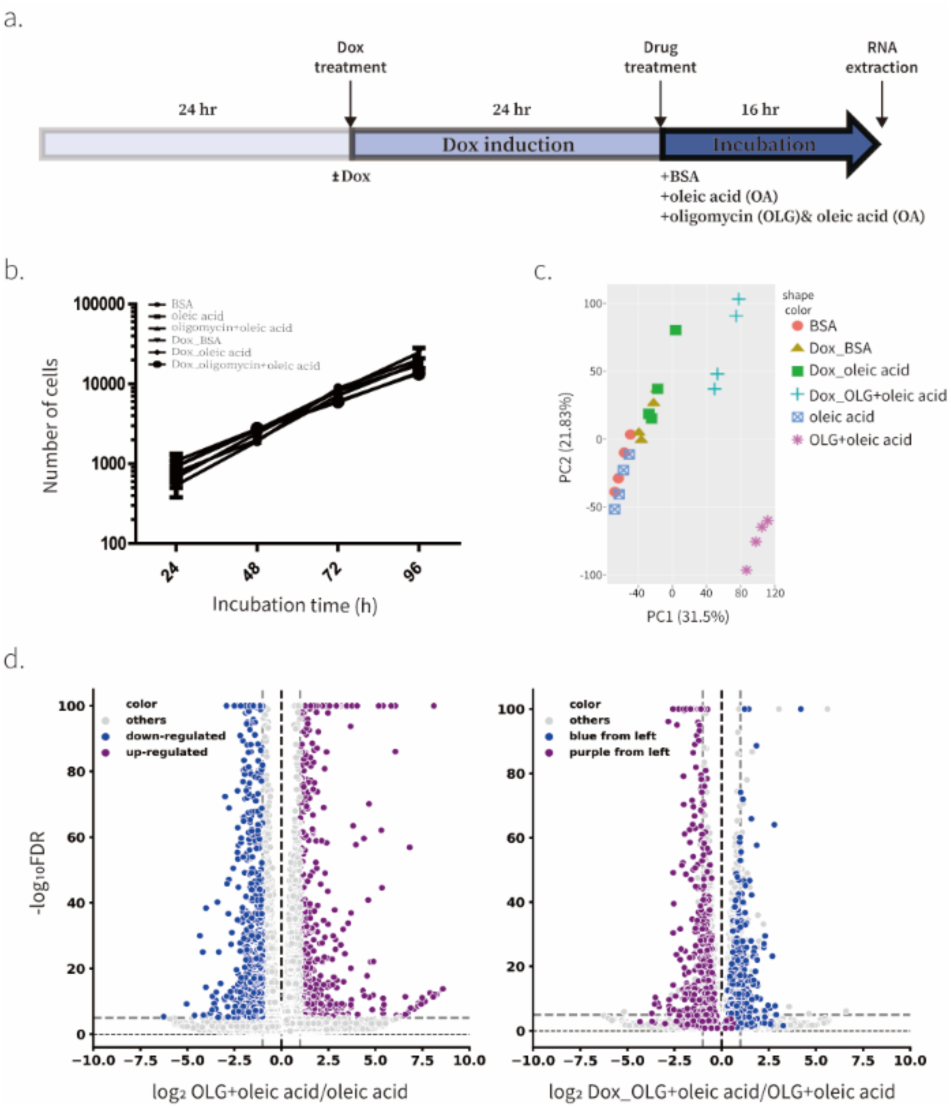
Transcriptional response to ATP synthase inhibition is driven by ΔΨm in C2C12 cells. **a**, A schematic diagram of RNA-seq experiment. **b,** Effect of various conditions included in RNA-seq on proliferation of C2C12 cells in the presence of 1 mM pyruvate. Means ± SEM, n=2 biological replicates. **c**, PCA of gene expression levels derived from RNA-seq in C2C12 treated with various combinations of doxycycline, BSA, oleic acid, and oligomycin for 16 hr. n=4 biological replicates. **d**, Fold-change (x-axis) and statistical significance (y-axis) of differential expression of genes focused on groups of genes responding to 16 hr oligomycin treatment. Genes showing differential expression in response to oligomycin-treatment (left) were traced in UCP1 overexpressed condition in the presence of oligomycin (right).

We next investigated whether the same genes were reciprocally regulated by oligomycin treatment and UCP1 expression in the presence of oligomycin. We labeled significantly upregulated and downregulated genes under the oligomycin-treated condition using two distinct colors: purple and blue, respectively (Fig. 5d). Strikingly, most downregulated genes (blue) were upregulated upon UCP1 expression in the context of oligomycin-treated conditions, while upregulated genes (purple) were downregulated (Fig. 5d). These transcriptional profiling results are consistent with our observations that UCP1 rescues oligomycin-induced ISR and suggests that ΔΨm changes drive most of the transcriptional changes in response to ATP Synthase inhibition.

## DISCUSSION

In this study, we validated UCP1 as a genetically encoded tool for direct manipulation of ΔΨm in live cells and showed that it is a more specific manipulator of ΔΨm than chemical uncouplers. We used UCP1 to show that increased ΔΨm drives ISR in response to ATP synthase dysfunction. We believe that this tool will complement chemical uncouplers and optogenetic tools by enabling the scientific community to study the regulation and physiological roles of changes in ΔΨm in living cells.

Most of our knowledge about the role of ΔΨm in cellular physiology stems from experiments using chemical uncouplers. Previous studies showed that treatment of chemical uncouplers reverses/prevents diet-induced obesity, insulin resistance, neurodegenerative and diabetic phenotypes in various model systems^35–41^. These observations suggested the possibility of targeting ΔΨm as a pharmacological therapy to achieve various beneficial effects. Our results suggest that chemical uncoupler have significant side effects independent of ΔΨm, such as inhibition of cell proliferation, that are observed at the same concentrations of uncpuplers that lead to maximal uncoupled respiration (Fig. 2f,g). Therefore, we suggest that caution should be exercised when attributing the effect of uncouplers to lowering of ΔΨm and that UCP1 should be used to verify key observations made with uncouplers.

We observed that UCP1 activity requires physiological levels of free fatty acids in the media, as expected since free fatty acids are required in catalytic amounts to stimulate UCP1 activity. This observation presents an advantage as the acute addition of free fatty acid to the media can be used to acutely manipulate ΔΨm in the presence of exogenous UCP1 in cells. At the same time, the concentrations of free fatty acids in mammalian organisms are similar or higher compared to the concentration that we used in the experiments to activate UCP1, which means that UCP1 should work in vivo without additional treatements^22,23^.

Previous studies have demonstrated that oligomycin, an ATP synthase inhibitor, induces a range of complex changes in metabolism, triggering the ISR^34,42^. These changes include NADH/NAD^+^ decrease, ATP depletion, and hyperpolarization of ΔΨm, which made it difficult to elucidate the bioenergetic driver of oligomycin-driven ISR^43–45^. Our genetic reagent allows for specific manipulation of ΔΨm without impeding cell proliferation, providing a unique advantage for studying the molecular mechanisms of ΔΨm-driven metabolic changes. In addition, this reagent holds significant promise across various research fields, particularly in investigating mitochondrial dysfunction associated with diseases linked to genetic mutations in ATP synthase subunits, including Leigh Syndrome, cerebellar atrophy, and encephalopathy^46–48^. Our study highlights the efficacy of directly manipulating ΔΨm in ameliorating oligomycin-induced metabolic changes, including ISR, offering insights into underlying mechanisms and potential therapeutic strategies for these disorders.

The increasing body of evidence underscores the importance of ΔΨm as a crucial parameter in the regulation of metabolism. Despite decades of studies employing chemical uncouplers to investigate the role of ΔΨm, limitations have become apparent due to their low specificity, leading to off-target effects on cellular metabolism. Recognizing this constraint, we aimed to develop a novel reagent devoid of such off-target effects. By validating UCP1 as a selective genetic tool for manipulating ΔΨm, we anticipate that this reagent will offer a valuable alternative to chemical uncouplers.

---

**Extended data Figure 1.**
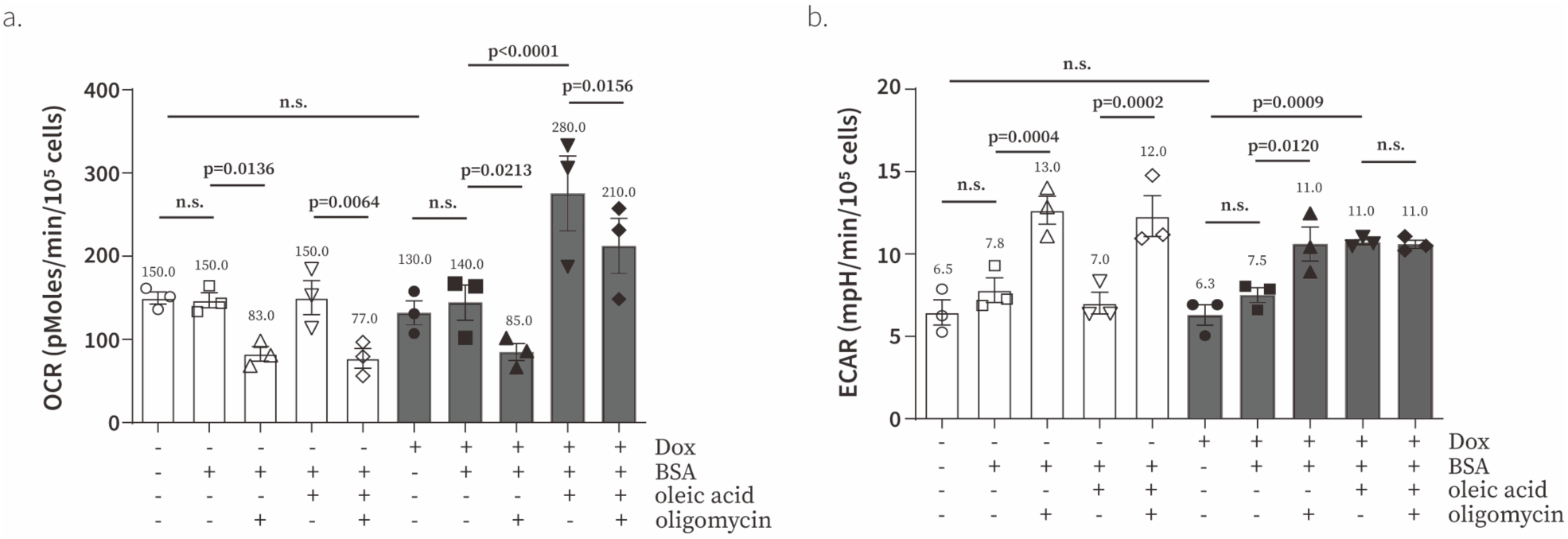
Effect of fatty acid or doxycycline on respiration and glycolysis in UCP1-expressing C2C12 cells. **a,b,** Effect of 51 μM BSA, 300 μM oleic acid, doxycycline (300 ng/ml), and 1 μM oligomycin in UCP1-expressing C2C12 cells were observed. Oxygen consumption rate (**a**) and extracellular acidification rate (**b**) were measured with an XFe24 analyzer. Means ± SEM, n=3 independent experiments. *P* values were determined with two-way ANOVA followed by Bonferroni’s multiple comparisons test in Graphpad (*n.s.*, not significant).

**Extended data Figure 2.**
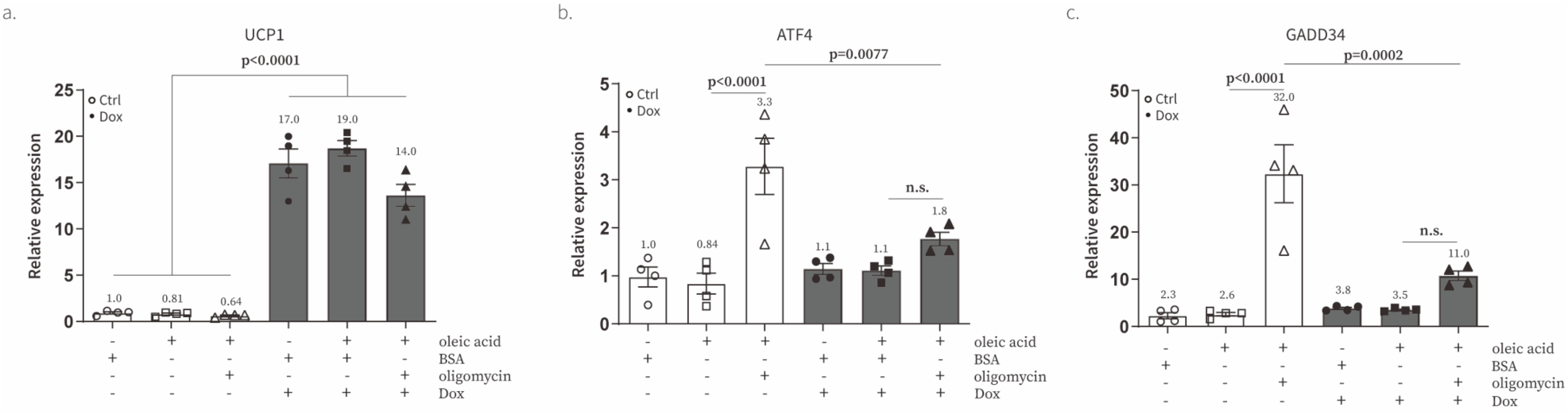
UCP1 rescues the effect of oligomycin on the transcription of ATF4 and downstream genes. **a-c**, qPCR of UCP1 (**a**), ATF4 (**b**), and GADD34 (**c**) following 16 hr treatment of combinations of doxycycline, 51 μM BSA, 300 μM oleic acid, and 1 μM oligomycin in the presence of pyruvate. Data is presented as fold-change to the condition with 51 μM BSA (BSA). Means ± SEM, n=4 biological replicates.

**Extended data Figure 3.**
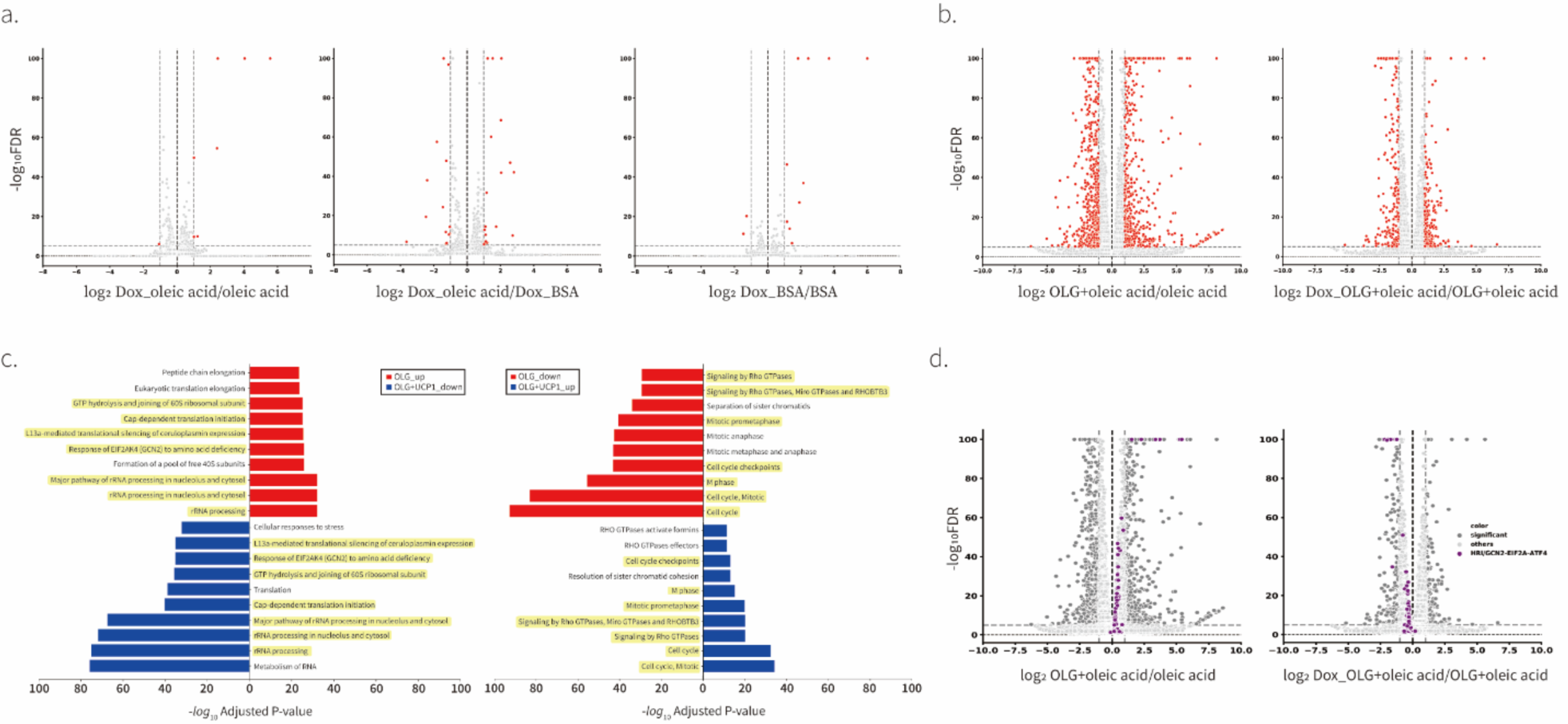
Validation of RNA-seq. **a-b**, Fold-change (x-axis) and statistical significance (y-axis) of differential expression of genes following 16 hr treatment of different combinations of doxycycline, BSA, oleic acid, and oligomycin. N=4 biological replicates. **c**, Gene ontology enrichment analysis of three different conditions. To determine the effect of oligomycin on the profile of transcription, analysis was performed to compare the data of oleic acid-treated to oleic acid with oligomycin-treated conditions. To determine the effect of UCP1 on oligomycin-driven changes in transcription, analysis was performed to compare the data of oleic acid with oligomycin-treated conditions in the absence and presence of doxycycline. The most significantly changed reactomes (top 10) are shown in the figure. Gene profiles of reactomes whose expression levels were reversed by UCP1 expression were shown with yellow highlights. N=4 biological replicates. **d**, Volcano plot relating a group of genes in EIF2α-ATF4 stress signaling under the condition of oligomycin treatment and UCP1 overexpression under the same oligomycin treatment. n=4 biological replicates.

## Acknowledgements

This work was supported by the National Institutes of Health grant DP2 GM132933 (D.V.T.). We thank the members of the Titov Lab for their valuable comments on the manuscript.

## Author contributions

Conceptualization: MC, DVT

Methodology: MC, DVT

Investigation: MC, DVT

Data Curation: MC, DVT

Visualization: MC, DVT

Funding acquisition: DVT

Writing – original draft: MC, DVT

Writing – review & editing: MC, DVT

## Data and materials availability

Vectors used in this study will be deposited on Addgene by the time of publication.

## Materials and Methods

### Cell lines and cell culture

Formulations of cell culture media used in this study are listed in Table S1. C2C12 cells were purchased from ATCC (CRL-1772) and were cultured in Dulbecco’s modified Eagle’s medium [low glucose DMEM without pyruvate (US Biological, D9802), 3.7 g/L NaHCO3, 10% FBS (Life Technologies, 10437028)]. Lentiviral-infected C2C12 cells were cultured in DMEM [low glucose DMEM without pyruvate (US Biological, D9802), 3.7 g/L NaHCO3, 10% FBS (Life Technologies, 10437028)] supplemented with 500 μg/ml geneticin (Gibco, 10131-035) and 10 μg/ml puromycin (Thermo Scientific, A11138-03). All of the experiments were performed in the absence of geneticin and puromycin.

All experiments were performed with cells before passage 7 for consistent phenotype. Cell lines were tested for mycoplasma contamination once every two months using the Mycotest kit (InvivoGen, rep-mys-10).

### Preparation of DNA constructs

Cloning of UCP1, mitoLbNOX, and LbNOX into pLVX-TRE3G vector

Mouse UCP1 gene and human codon-optimized mitoLbNOX and LbNOX (addgene #74448 and #75285) were cloned into pLVX-TRE3G vector (Clontech, CA) using Gibson assembly. All constructs also included strong Kozak sequence (<u>GCCACCATGGGG</u>) at the upstream of the start codon.

### Lentivirus production

1.3 million HEK293T cells were seeded in a 6 cm dish in 5 ml of DMEM [high glucose DMEM (Gibco, 12800), 3.7 g/L NaHCO3, 10% FBS (Life Technologies, 10437028)]. After 18 hr, cells were transfected with 300 μl transfection mixture. The transfection mixture contained 6 μl of Lipofectamine 2000 (Invitrogen, 11668-027), 1400 ng of psPAX2 (Addgene plasmid #12260), 140 ng of pMD2.G Addgene plasmid #12259), 1400 ng of pLVX-TRE3G vector of interest (including pLVX-TRE3G-Luc control vector, expressing Luciferase, purchased from Clontech) and Opti-MEM medium (Gibco, 31985062) up to 300 μl. To make the transfection mixture, 150 μl solutions of Lipofectamine 2000 and DNA mixtures were prepared separately, and DNA solution was added dropwise to the Lipofectamine 2000 solution. The mixture was incubated at room temperature for 30 min before adding it to cells. Two days after transfection, medium was collected, filtered through 0.45 μm filter. Filtered medium was centrifuged at 500 x *g* for 5 min to pellet remaining cells and the supernatant was used infect cells. To prepare concentrated lentivirus, 1 volume of Lenti-X concentrator (Clontech, 631231) was added to 3 volumes of clarified-lentivirus containing medium. The mixture was incubated at 4 °C after gentle inversion to mix well. After 18 hr, the mixture was centrifuged at 1500 x *g* at 4 °C for 45 min and completely remove supernatant. Add 1/10 to 1/100 of the original volume using complete DMEM and used to infect cells.

### Generation of stable cells using lentiviral infection

Twenty thousand C2C12 cells were seeded in 2 ml of DMEM [low glucose DMEM without pyruvate (US Biological, D9802), 3.7 g/L NaHCO3, 10% FBS (Life Technologies, 10437028)] with 10 μg/ml of polybrene (Sigma, TR-1003) per well in a 6-well plate and various volumes of DMEM medium containing lentivirus was added per well. Medium was replaced with 2 ml of DMEM [low glucose DMEM without pyruvate (US Biological, D9802), 3.7 g/L NaHCO3, 10% FBS (Life Technologies, 10437028)] ± 500 μg/ml geneticin (Gibco, 10131-035) ± 1 μg/ml puromycin (Thermo Scientific, A11138-03). Stable cells were selected for at least a week before performing experiments and were cultured in the presence of the indicated concentrations of antibiotics.

### Preparation of BSA-conjugated oleic acid

BSA-oleic acid conjugate was prepared based on the BSA-palmitate conjugate preparation protocol (SeahorseBioscience) with minor modifications. Fatty acid free albumin (Sigma, 10775835001) and sodium oleate without methyl-β-cyclodextrin (Sigma, O7501) were used and 1:6 ratio of BSA to oleic acid were used to prepare conjugate. BSA (1.02 mM) and oleic acid (7.5 mM) were dissolved in 150 mM NaCl solution at ∼70 °C and < 37 °C, respectively. The mixture of solutions (BSA:oleic acid:NaCl solution=5:4:1) was stirred for 1 hr at 37 °C and adjusted pH to 7.4 by adding 1 M NaOH. BSA solution without oleic acid was also prepared by 2-fold dilution of BSA solution with 150 mM NaCl solution and final pH was adjusted to 7.4 by adding 1 M NaOH. Aliquots were stored at −80 °C before use.

### Oxygen consumption rate (OCR) and Extracellular acidification rate (ECAR) of cell lines expressing UCP1 (Fig. 1D and extended data Fig. 1)

Oxygen consumption rates (OCR) and Extracellular acidification rate (ECAR) of C2C12 cells expressing UCP1 under the control of the doxycycline-inducible promoter (TRE3G) were measured with Seahorse XFe24 analyzer (Agilent). Cells were seeded at 4 x 10^4^ cells per well in XFe24 cell culture microplates (Agilent, 102340-100) in 10 ml of DMEM [high glucose DMEM with pyruvate (Gibco, 12800), 3.7 g/L NaHCO3, 10% FBS (Life Technologies, 10437028)] and were incubated at 37 °C in 5% CO_2_ incubator. Medium was replaced the next day with 1 ml of fresh medium per well and doxycycline (final concentration of 300 ng/ml, prepared in water) or water was added to the corresponding wells to induce protein expression. After 24 hr, cells were washed with 1 ml of the assay medium [high glucose with pyruvate (Gibco, 12800), 10% FBS (Life Technologies, 10437028) and 25 mM HEPES-KOH, pH 7.4]. Then medium was replaced with 0.5 ml of the same assay medium and plates were placed in the Seahorse XFe24 analyzer for measurement of OCR. Each measurement was performed over 4 min after a 2-min mix and a 2-min wait period. Basal measurements were collected 3 times and 3 measurements were collected after injection of oleic acid (final concentration of 300 μM and injection volume of 56 μl), 3 measurements were collected after injection of oligomycin (final concentration of 1 μM and injection volume of 62 μl), 3 measurements were collected after injection of FCCP (final concentration of 5 μM and injection volume of 69 μl), followed by 3 measurements after addition of antimycin A (final concentration of 1 μM and injection volume of 75 μl). Both Oxygen consumption rates (OCR) and Extracellular acidification rates (ECAR) were normalized by the cell number.

### Oxygen consumption rate (OCR) under various concentrations of uncouplers (FCCP or Bam15) (Fig. 2c)

Oxygen consumption rates (OCR) of wild-type C2C12 under various concentrations of uncouplers were measured with Seahorse XFe24 analyzer (Agilent). Experiments were conducted under the same process as explained above. Basal measurements were collected 3 times and 12 measurements were collected after injection of uncouplers (FCCP or Bam15, final concentrations of 0, 0.625, 1.25, 2.5, 5, 10, and 20 μM. injection volume of 56 μl), 5 measurements were collected after injection of oligomycin (final concentration of 1 μM and injection volume of 62 μl), followed by 3 measurements after addition of antimycin A (final concentration of 1 μM and injection volume of 69 μl). Oxygen consumption rates (OCR) were normalized by the cell number.

### Western blot

Cells were lysed using 2X SDS-PAGE sample buffer (24 mM Tris-HCl, pH 6.8, 10% glycerol, 0.8% SDS, and 286 mM β-mercaptoethanol). Samples were subjected to SDS-PAGE on 4-20% Mini-PROTEAM^R^ TGX^TM^ gels (Bio-Rad, 4561096) and transferred to a nitrocellulose membrane (Bio-Rad, 1704159). The primary antibodies used in this study were: rabbit anti-ATF4 (Cell Signaling Technologeis, 11815s, 1:625), rabbit anti-β-actin (Cell Signaling Technologies, 4970S, 1:2500), rabbit anti-P-EIF2α (Cell Signaling Technologies, 3398S, 1:1000), mouse anti-EIF2α (Fisher Scientific, AHO0802, 1:1000), rabbit anti-β-Tubulin (Thermo Fisher, 32-2600, 1:2500), rabbit anti-UCP1 (Abcam, ab10983, 1:2500), rabbit anti-Flag (Cell Signaling Technologies, 2368S, 1:5000), rabbit anti-PKM2 (Cell Signaling Technologies, 4053S, 1:1000), rabbit anti-Tom20 (Cell Signaling Technologies, 42406S, 1:2500). Blots were incubated with anti-rabbit IgG (Cell Signaling Technologies, 7076S, 1:10000) or anti-mouse IgG (Cell Signaling Technologies, 7074S, 1:10000) and imaged using an analyticjena imaging system. Digital images were processed and quantified using ImageJ software.

### Drug treatment_ISR assay (Fig. 3 and extended data Fig. 4, 5)

C2C12 cells and derived cell lines were seeded with 0.1 x 10^6^ cells per well in 6-well cell culture plates. After 24 hr, doxycycline (final concentration of 300 ng/ml, prepared in water) or water was added to the corresponding wells to induce protein expression. After 24 hr, medium was replaced with the DMEM medium [high glucose with pyruvate (Gibco, 12800), 3.7 g/L NaHCO3, 10% FBS (Life Technologies, 10437028)] containing final drug concentrations. Cells were incubated at 37 °C with the following mitochondrial toxins for 16 hr: 1 μM oligomycin (Signa-Aldrich, 75351), 5 μM FCCP (TOCRIS, 45310), 1 μM antimycin A (Sigma-Aldrich, A8674), 1 μM rotenone (Sigma-Aldrich, R8875), 50 μgml^-1^ doxycycline (Sigma-Aldrich, D9891). After 16 h, medium was aspirated, and cells were washed once with 2 ml of ice-cold PBS. Cells were lysed with 150 μl of 2X SDS-PAGE sample buffer (24 mM Tris-HCl, pH 6.8, 10% glycerol, 0.8% SDS, and 286 mM β-mercaptoethanol).

### Cell proliferation assays

#### Rescure of oligomycin-induced inhibition of cell proliferation and the effect of pyruvate on oligomycin-induced inhibition (Extended data Fig. 2a)

Two hundred UCP1-expressing C2C12 cells were seeded in 200 μl of DMEM [high glucose DMEM with pyruvate (Gibco, 12800), 3.7 g/L NaHCO3, 10% FBS (Life Technologies, 10437028)] per well in a black 96-well plate with a clear bottom (Corning, 3904). Twenty-four hours after seeding, medium was exchanged to DMEM [high glucose DMEM with pyruvate (Gibco, 12800), 3.7 g/L NaHCO3, 10% FBS (Life Technologies, 10437028)] ± 1 μM oligomycin ± 300 ng/ml doxycycline ± 300 mM oleic acid or DMEM [low glucose DMEM without pyruvate (US Biological, D9802). 1.8 g/L glucose, 3.7 g/L NaHCO3, 10% FBS (Life Technologies, 10437028)] ± 1 μM oligomycin (or 1 μM piericidin) ± 300 ng/ml doxycycline ± 300 mM oleic acid. After 0, 1, 2, and 3 days, medium was aspirated, and cells were fixed by adding 100 μl of 4% paraformaldehyde in PBS and incubating at 37 °C for 15 min. Paraformaldehyde solution was aspirated and cells were stained with 100 ul of 1 μg/ml Hoechst 33345 (Invitrogen, H21492) in PBS. Plates were covered with sealing aluminum foil (VWR, 60941-112) and stored at 4 °C before counting cells in each well with Cytation1 (see imaging protocol below).

#### Rescue of oligomycin (or piericidin)-induced inhibition of cell proliferation (Extended data Fig. 2b)

Four hundred UCP1-expressing C2C12 cells were seeded in 200 μl of DMEM [high glucose DMEM with pyruvate (Gibco, 12800), 3.7 g/L NaHCO3, 10% FBS (Life Technologies, 10437028)] per well in a black 96-well plate with a clear bottom (Corning, 3904). Twenty-four hours after seeding, medium was exchanged to DMEM [high glucose DMEM with pyruvate (Gibco, 12800), 3.7 g/L NaHCO3, 10% FBS (Life Technologies, 10437028)] ± 1 μM oligomycin (or 1 μM piericidin) ± 300 ng/ml doxycycline ± 300 mM oleic acid. After 0, 1, 2, and 3 days, medium was aspirated, and cells were fixed by adding 100 μl of 4% paraformaldehyde in PBS and incubating at 37 °C for 15 min. Paraformaldehyde solution was aspirated and cells were stained with 100 ul of 1 μg/ml Hoechst 33345 (Invitrogen, H21492) in PBS. Plates were covered with sealing aluminum foil (VWR, 60941-112) and stored at 4 °C before counting cells in each well with Cytation1 (see imaging protocol below).

#### Effect of doxycycline, oleic acid, and UCP1 on cell proliferation (Fig. 2e)

Four hundred wild-type or UCP1-expressing C2C12 cells were seeded in 200 μl of DMEM [high glucose DMEM with pyruvate (Gibco, 12800), 3.7 g/L NaHCO3, 10% FBS (Life Technologies, 10437028)] per well in a black 96-well plate with a clear bottom (Corning, 3904). Twenty-four hours after seeding, medium was exchanged to DMEM [high glucose DMEM with pyruvate (Gibco, 12800), 3.7 g/L NaHCO3, 10% FBS (Life Technologies, 10437028)] ± 300 ng/ml doxycycline ± 300 mM oleic acid. After 0, 1, 2, and 3 days, medium was aspirated, and cells were fixed by adding 100 μl of 4% paraformaldehyde in PBS and incubating at 37 °C for 15 min. Paraformaldehyde solution was aspirated and cells were stained with 100 ul of 1 μg/ml Hoechst 33345 (Invitrogen, H21492) in PBS. Plates were covered with sealing aluminum foil (VWR, 60941-112) and stored at 4 °C before counting cells in each well with Cytation1 (see imaging protocol below).

#### Nuclei counting using Cytation1

Images of 96-well plates with fixed cells stained with Hoechst 33345 were collected using Cytation1 (Agilent). Twenty images were taken to cover the whole well. Images were analyzed and nuclei number per well was counted using Gen5^TM^ version 3.05 (Agilent). The cell counting method had a linear range from 100 to 30000 cells per well as determined by counting plates with known number of cells seeded 12 hr prior to fixation.

#### Measurement of mitochondrial membrane potential with fluorescence microscope using TMRM (Fig. 2a,b)

Ten thousand wild-type or UCP1-expressing C2C12 cells were seeded per well of 24-well glass bottom plate (MatTek, P24G1.513F) in 1 ml of DMEM [high glucose DMEM with pyruvate (Gibco, 12800), 3.7 g/L NaHCO3, 10% FBS (Life Technologies, 10437028)]. Twenty-four hours after seeding, 1/20 volume of doxycycline (final concentration of 300 ng/ml) or same volume of water was added. Twenty-four hours after doxycycline addition, medium was exchanged to 1 ml of DMEM [low glucose DMEM without pyruvate (US Biological, D9802), 1.8 g/L glucose, 10% dialyzed FBS (ThermoFisher, 26400044) with 25 mM HEPES-KOH, pH 7.4] with the indicated combination of 10 nM TMRM (Invitrogen, T668) and 1 μg/ml Hoechst 33345. UCP1-expressing cells were treated with 50 μM Cyclodextrin conjugated oleic acid (Sigma, O1257) and one of the following drugs – 1 μM oligomycin or 5 μM FCCP. Wild-type C2C12 cells were treated with various concentrations of uncouplers (FCCP, 0-8 μM; Bam15, 0-20 μM). One hour incubation at 37 oC without CO_2_, fluorescence image was captured using Cytation1. Fluorescence intensity was determined by ImageJ v1.53 and intensities of TMRM were normalized by intensity of Hoechst 33345. Relative fluorescence intensities were shown compared to the control conditions (first lane of each bar graph).

#### Determination of UCP1 localization using cell fractionation experiment (Fig. 1c)

One million UCP1-expressing C2C12 cells were seeded in 10-cm cell culture dish in 10 ml of DMEM [high glucose DMEM with pyruvate (Gibco, 12800), 3.7 g/L NaHCO3, 10% FBS (Life Technologies, 10437028)] and doxycycline (300 ng/ml final concentration) or same volume of water was added 24 hr after seeding cells. Twenty-four hours after doxycycline addition, medium was removed, and cells were washed once with 5 ml of ice-cold PBS. One ml of ice-cold fractionation buffer (0.2 M sucrose, 10 mM Tris-HCl, pH 7.4, 1 mM EGTA, pH 7.4, with 1X protease inhibitor (Roche, 11836170001)) was added to cells and cells were harvested using a cell scraper. Cells were centrifuged at 200 x *g* for 5 min and supernatant was completely removed. Cells were resuspended in 800 μl of fractionation buffer and disrupted using Teflon homogenizer (DWK Life Sciences, 3432S10) for 10 times. In order to remove intact cells, debris, cell membranes, and nuclei from the supernatant containing cytoplasm, mitochondria, and other small membrane-bound organelles, cell lysate was centrifuged at 1000 x *g* for 5 min at 4 °C. The supernatant was transferred to fresh Eppendorf tubes and centrifuged for 10 min at 10000 x *g* at 4 °C. The resulted supernatant was the cytosolic fraction and pellet containing mitochondria was washed after removing supernatant (cytosolic fraction) by resuspending in 200 μl of ice-cold fractionation buffer and centrifuged at 10000 x *g* for 2 min to remove residual cytosolic components (repeat 2 times). Mitochondrial pellet was resuspended in 50 ul of SDS-PAGE sample buffer (24 mM Tris-HCl, pH 6.8, 10% glycerol, 0.8% SDS, and 286 mM β-mercaptoethanol) and stored at 4 °C before use. Antibodies against PKM2 (Cell Signaling Technologies, 4053S), UCP1 (Abcam, ab10983), and Tom20 (Cell Signaling Technologies, 42406S) were used.

#### Determination of UCP1 localization using immunofluorescence (Fig. 1d)

One hundred thousand UCP1-expressing C2C12 cells were seeded on fibronectin-coated glass coverslips in 12-well plates in 1 ml of DMEM [high glucose with pyruvate (Gibco, 12800), 3.7 g/L NaHCO3, 10% FBS (Life Technologies, 10437028)]. After 24 hr of seeding, doxycycline (final concentration of 300 ng/ml) or equal volume of water was added. After 24 hr, medium was completely aspirated and cells were fixed using 4% paraformaldehyde (Fisher Scientific, 50980487) for 15 min at room temperature. Cells were then rinsed two times with 2 ml of PBS, permeabilized with 0.1% (w/v) Triton-X in PBS for 20 min at room temperature, and rinsed again three times with PBS. Primary antibodies (diluted 1:300) were diluted into 5% normal donkey serum (Jackson ImmunoResearch, 017-000-121) and incubated overnight at 4 °C. The following day, coverslips were rinsed three times with DPBS and then labeled with fluorescently-conjugated secondary antibodies (diluted 1:400 in 5% normal donkey serum in DPBS) for 1 hr at room temperature, protected from light. Coverslips were rinsed with PBS four times and mounted on glass slides using VECTASHIELD Antifade Mounting Medium with DAPI (Vector Laboratories, H-1000). All confocal microscopy was performed using a spinning-disk confocal system built on a Nikon Eclipse Ti microscope.

#### Cell treatment for RNA isolation (Fig. 4a)

Doxycycline-inducible C2C12 cells were seeded for most treatments at 0.1X10^6^ cells per well in 6-well plates with 2 ml of DMEM [high glucose DMEM with pyruvate (Gibco, 12800), 3.7 g/L NaHCO3, 10% FBS (Life Technologies, 10437028)] without selection antibiotics. After 24 hr of seeding, 100 μl of doxycycline (final concentration of 300 ng/ml, prepared in water) or water was added to the corresponding wells to induce protein expression. After 24 hr, various treatments were started with complete replacement of the media to a final volume of 2 ml/well (still containing 300 ng/ml doxycycline or water). BSA (Sigma, 10775835001) was used at 51 μM, oleic acid (Sigma, O1383) at 300 μM, and oligomycin (Sigma, 75351) at 1 μM. Treatments intended for RNA isolation lasted 16 hr. At the end of the treatment, media were completely aspirated, and cells were lysed in 350 μl/well lysis buffer from the Aurum^TM^ total RNA Mini Kit (Biorad, 7326820). The lysates were immediately frozen at −80 °C until RNA extraction. RNA isolation was performed using the Aurum^TM^ total RNA Mini Kit following the manufacturer’s instructions.

#### RNA sequencing

The concentration and quality of RNA intended for sequencing was assayed using nanodrop one (Thermo Scientific). RNA samples (1000 ng/μl for each conditions) were sent to Novogene Corporation Inc. for RNA-sequencing. RNA-seq libraries were prepared using the NEBNext Ultra II RNA Library prep kit from Illumina according to the manufacturer’s instructions and sequenced as PE150 reads on an Illumina NovaSeq 6000. The sequencing data will be deposited in the NCBI Gene Expression Omnibus (GEO) database by the time of publication.

#### qPCR (Extended data Fig. 3)

Isolated RNA (1000 ng) was mixed with reverse transcription master mix and reverse transcribed based on following steps (5 min at 25 °C, 20 min at 45 °C, 1 min at 95 °C) using iScript Reverse Transcription Supermix for RT-qRCR (Biorad, 1708840). The resulting cDNA was diluted by adding 24X volume of DPEC water subjected to qPCR using SsoAdvanced^TM^ Universal SYBR Green Supermix (Biorad, 1725271) on StepOnePlus Real-Time PCR (Applied Biosystems).

Isolated RNA was annealed to random primers (Thermo Fisher Scientific; 48190–011) for 5 min at 70C, then reverse transcribed for 1 hr at 37 °C using M-MLV reverse transcriptase (Promega, Madison, WI; M1705) in the presence of RNase inhibitor (Thermo Fisher Scientific; 10777–019). The resulting cDNA was subjected to qPCR using TaqMan gene expression master mix (Thermo Fisher Scientific; 4369514) and TaqMan gene expression probes (see Key Resources Table for details) on a CFX96 instrument (Bio-Rad, Hercules, CA). Raw amplification cycle data was produced by the accompanying analysis software using default parameters. Cycle differences between tested conditions and the baseline condition were normalized against the reference gene Ubr3 (for mouse samples) or TBP (for human samples), yielding ΔΔC_t_. Fold-changes were calculated as 2^-1ΔΔCt^, but statistical testing was performed on the underlying ΔΔC_t_ values.

